# Parasites Acquired Beta Satellite DNAs from Hominid Hosts via Horizontal Gene Transfer

**DOI:** 10.1101/589531

**Authors:** Jiawen Yang, Yiting Zhou, Guangwei Ma, Xueyan Zhang, Yabin Guo

## Abstract

Beta satellite DNA (satDNA) sequences are repeated DNA elements located in primate centromeres and telomeres, and might play roles in genome stability and chromosome segregation. Beta satDNAs mainly exist in great apes. Previous studies suggested that beta satDNAs may originate in old world monkeys. In this study, we searched both GenBank and SRA database, and identified beta satDNA sequences from the genomic sequences of 22 species. The beta satDNA sequences found in Prosimian, Dermoptera and Scandentia indicated that the origin of beta satDNAs might be as early as 80 MYA. Strikingly, beta satDNA sequences were also found in a number of some species evolutionarily far from primates, including several endoparasites of human and other great apes, which could be the results of multiple horizontal gene transfer (HGT) events. The similar phylogenic profiles between beta satDNAs in the parasite genomes and the human genome indicates that the parasite beta satDNAs have undergone similar concerted evolution and play similar roles as the beta satDNAs in primates.

**Highlights:**

1. The ever largest scale analysis on beta satDNAs.
2. The origin of beta satDNAs was traced back to ∼80 MYA.
3. Mass existence of beta satDNAs in non-primate species was contributed by multiple HGT events.

## Introduction

The genomes of eukaryotes comprise large tracts of repeated sequences. More than half of the human genome are repetitive sequences (Giussani et al., 2012). Repetitive DNA sequences include satellite DNAs (satDNAs), minisatellite, microsatellite sequences and transposable elements (Charlesworth et al., 1994). Highly homogenized arrays of tandem repeats, known as satDNAs, are enriched in centromeric, pericentromeric, subtelomeric regions and interstitial positions (Yunis and Yasmineh, 1971). SatDNAs have recently been reconsidered to have various functions, such as playing roles in genome stability and chromosome segregation (Khost et al., 2017). Furthermore, they may even involve mammalian gene regulations (Tomilin, 2008). The changes in the copy number of repetitive sequences in genomic DNA are important causes of hereditary disorders (J. Rich et al., 2014; Mirkin, 2006, 2007). The insertion of 18 beta satellite units in the gene coding a transmembraine serine protease causes congenital and childhood onset autosomal recessive deafness (Scott et al., 2001). The global DNA hypomethylation frequently observed in cancers is mostly taken place at satDNAs (J. Rich et al., 2014). The loss of the BRCA1 tumor suppressor gene provokes satDNA derepression in breast and ovarian tumors in both mice and humans (Zhu et al., 2011). Facioscapulohumeral muscular dystrophy (FSHD), a autosomal dominant hereditary disease, is associated with the macrosatellite (D4Z4) and beta-satellite (4qA allele) DNA sequences (J. Rich et al., 2014).

Beta satDNAs were considered to be unique in the primates. The basic units of beta satDNAs are 68 bp long with a higher GC content. Beta satDNAs are also known as Sau3A sequences for the presence of a Sau3A restriction site within nearly every single unit (Meneveri et al., 1985). In the human genome, Beta satDNAs have been identified in the pericentromeric regions of chromosomes 1, 9, and Y (Meneveri et al., 1993), as well as in acrocentric chromosomes (13, 14, 15, 21 and 22) (Agresti et al., 1987) and in chromosome 19p12 (Eichler et al., 1998). Beta satDNAs were also described in great apes (human, chimpanzee, gorilla, and orangutan), lesser apes (gibbons) (Meneveri et al., 1995) and old world monkeys (Cardone et al., 2004). However, most of these studies were performed before next generation sequencing was utilized and the amount of sequences studied was relatively limited. Today, taking the advantage of high-throughput sequencing, many more satDNA sequences can be identified than ever for more comprehensive investigations.

In this study, we analyzed the distribution of beta satDNAs in 22 species and provide a landscape of beta satDNA evolution. The beta satDNAs were also found in non-primate species besides in primates, which could be the results of horizontal gene transfers (HGT).

## Results

### Beta satDNAs in the human genome

The current human genome assembly is not yet complete for the high degree of sequence homogeneity among many hundreds or thousands of copies of repeated sequences. Most human chromosome contigs contain few or no beta satDNA sequences except chromosome Y. The beta satDNA sequences on chromosome Y are clustered in three separated regions designated as Ya, Yb and Yc in this study, which were shown in Fig.1A. The sequences of the three regions were mapped and single units of beta satDNA sequences were extracted. After CLUSTALW alignments being performed, the phylogenetic relationships of these sequences were examined using RAxML software (Fig. 1B).

**Fig. 1.**
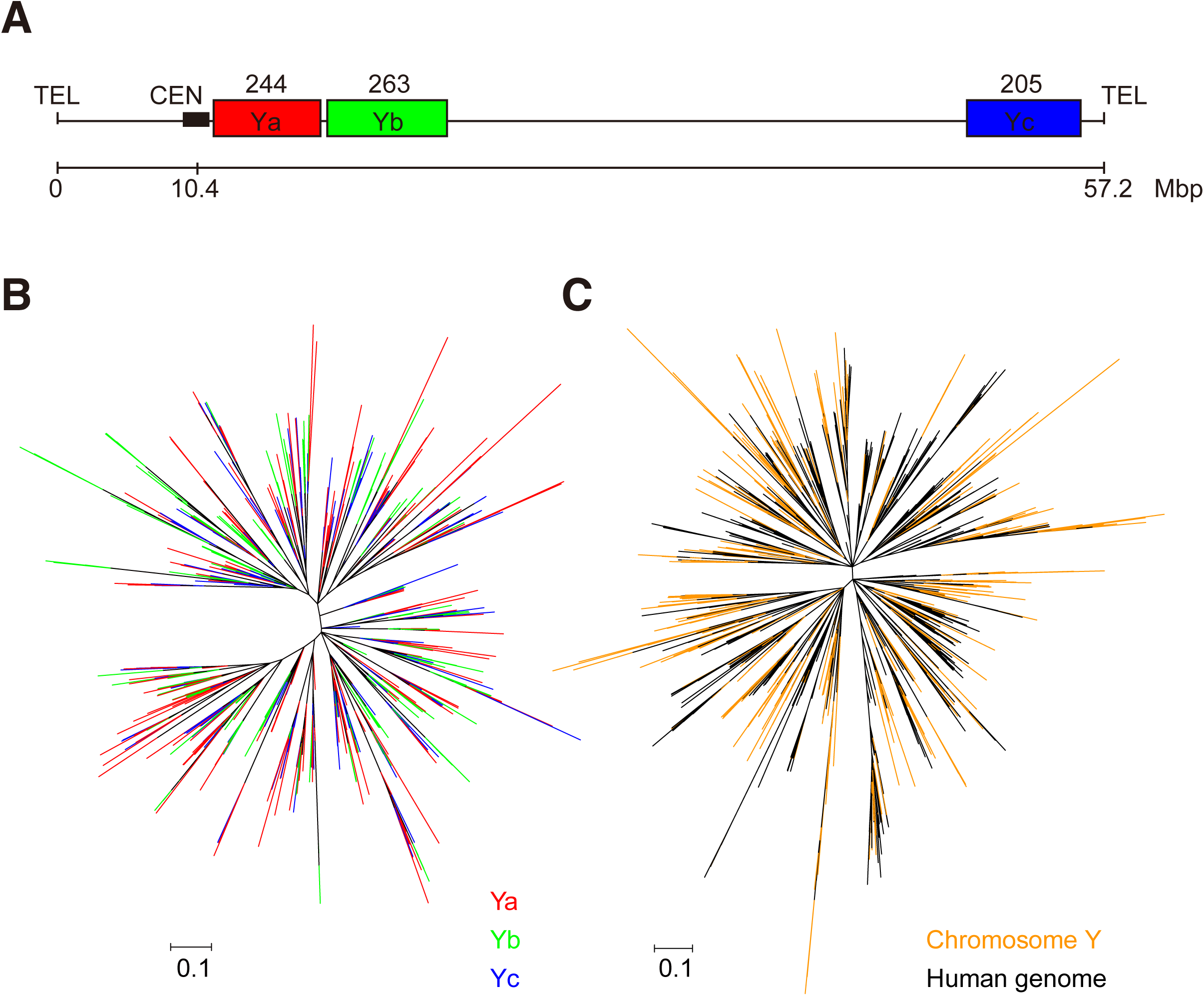
Distribution and diversity of beta satDNA sequences on human chromosome Y. A, Beta satDNA sequences on the chromosome Y. The beta satDNA copy numbers were labeled above the boxes. B, Phylogenetic tree of beta satDNAs on chromosome Y. Ya, Yb and Yc refer to beta satDNAs located at regions shown in A. C, Phylogenetic tree of beta satDNAs on chromosome Y (orange) and 1,000 copies beta satDNAs randomly picked from the human genome (black). The trees were constructed using RAxML with statistical support provided by bootstrapping over 1,000 replicates.

Obviously, the sequences from Ya, Yb and Yc are not phylogenetically separated. To see if the beta satDNA sequences on chromosome Y are different from those on other chromosomes, we randomly picked 1000 copies of non-repeated beta satDNA sequences from the human genome raw WGS (whole genome sequencing) data to build a phylogenetic tree together with the beta satDNA sequences on chromosome Y (Fig. 1C). This result shows that there is no regional or chromosomal specificity in human beta satDNA sequences, indicating that the evolution of beta satDNAs, similar to the higher-order alpha satDNAs, are extremely homogeneous within and between chromosomes, which may be a result of intrachromosomal and interchromosomal exchanges (Rudd et al., 2006). The beta satDNA sequences on chromosome Y have relatively higher diversity among the beta satDNA sequences in human genome, which is consistent with the previous report (Cardone et al., 2004).

### The distribution of beta satDNAs in different species and putative horizontal transfers

To analyze the diversity and evolutionary relationships of beta satDNAs in different species, we tried to collect as many sequences as possible. First, we searched all the available sequences in GenBank using NCBI-BLAST, and found a number of species possibly containing beta satDNAs. Since beta satDNAs are highly repeated and located at centromeres and telomeres, large number of beta satDNAs copies haven’t been assembled into chromosome contigs. The copy numbers of beta satDNAs found in assembled genomes are very limited. To obtain more sequence copies for better characterization, we directly searched beta satDNA sequences for the species found during the previous step within the raw WGS data in the NCBI SRA (Sequence Read Archive) database. Eventually, overall 215,239 beta satDNA sequences were identified in 22 species (Supplementary file 1, Table S1).

Besides in apes, beta satDNAs were found in old world monkeys, *Macaca fascicularis* and *Mandrillus leucophaeus*, and in new world monkeys, *Callithrix jacchus* (marmoset) and *Saimiri boliviensis* (squirrel monkey). We also found beta satellite sequences in *Carlito syrichta* (tarsier) and *Prolemur simus* (greater bamboo lemur), and even in *Galeopterus variegatus* (Sanda flying lemur) and *Tupaia belangeri* (tree shrew), which indicates the origin of beta satDNAs could be as early as 80 MYA. Interestingly, beta satDNAs were also found in genomes of some non-Primatomorpha species, including cattle, tape worms, roundworms and even plant.

The proportions of beta satDNAs in different species vary greatly. The hominid genomes have the highest proportions of beta satDNAs; the genomes of lesser apes and old world monkey have modest proportions, while other mammalian genomes have the least proportions. However, the proportions of beta satDNAs in *Spirometra erinaceieuropaei* (tapeworm) and *Enterobius vermicularis* (pinworm) are considerably high (Fig. 2A).

**Fig. 2.**
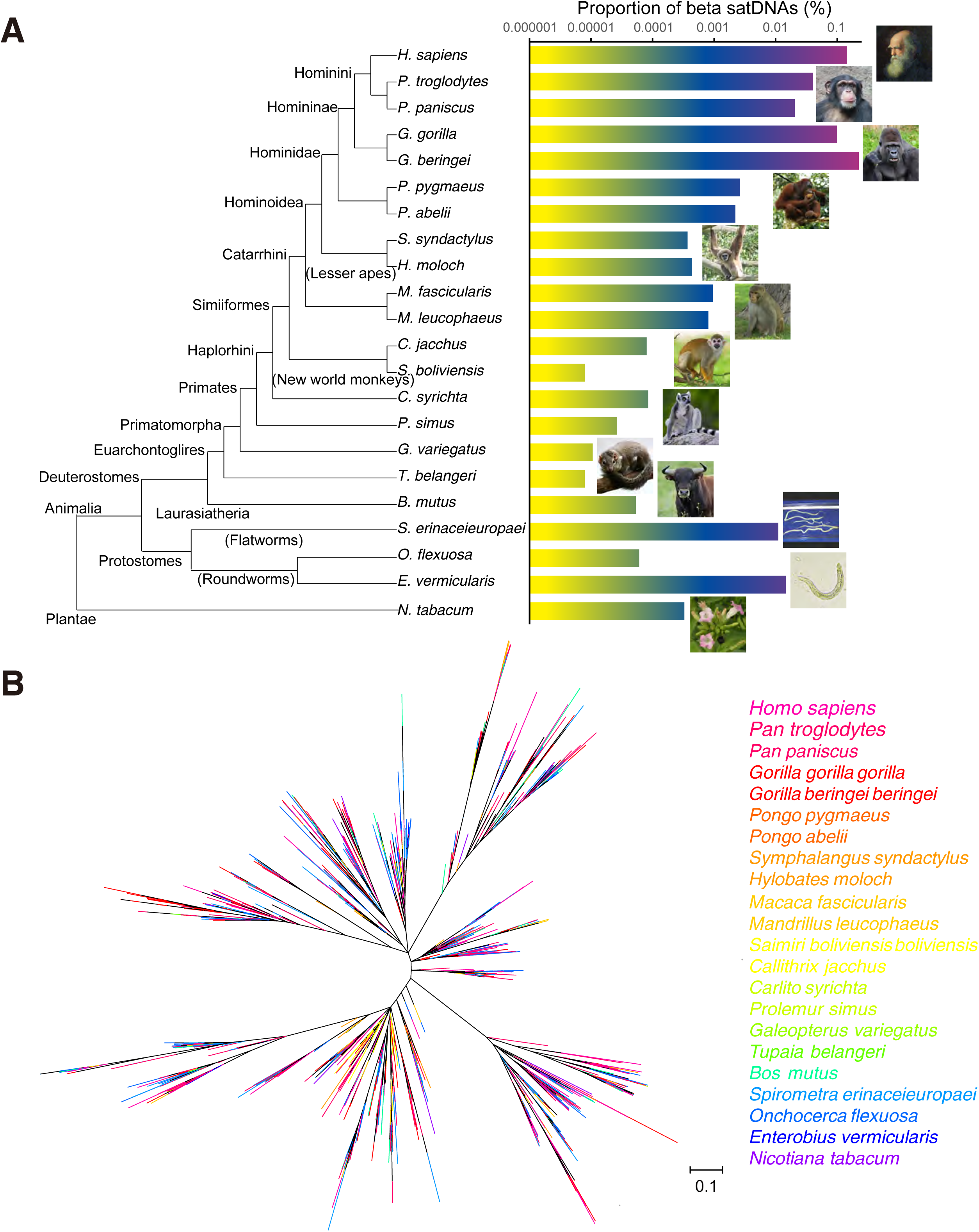
The distribution and diversity of beta satDNAs in different species. A, the proportions of beta satDNAs in different species were shown by the histogram at the right part. The evolutionary relationships of the species were shown at the left part. The lengths of the lines of the phylogenetic tree didn’t reflect the genetic distances between species. B, Phylogenetic tree of 2,830 beta satDNA sequences of different species was constructed using RAxML, with statistical support provided by bootstrapping over 1,000 replicates.

Considering the possibility that the DNA samples of other species could be contaminated by human DNA, we excluded all the beta satDNA sequences in the non-human WGS data, if they can also be found in the human WGS data, resulting in a new dataset with higher stringency (Table S1). The sequence distribution of the high-stringency dataset is similar to the original one, except that the sequence numbers in new world monkey, Dermoptera and Scandentia changed to zero, but the sequences in tarsier and lemur were still retained.

To analysis the diversity and evolution of the beta satDNA sequences, we examined the phylogenetic relationships with 2,830 beta satDNA sequences from the 22 species. The sequences taken within each species were all different. For the species with more than 300 different sequences, 300 sequences were randomly picked, while for the rest of the species, all available sequences were chosen. Thus, a complex phylogenetic tree with a mixed-clade group of 2,830 individuals of the 22 species was built (Fig. 2B). Basically, beta satDNA sequences from each of the 22 species can be found in any of the main clusters of the tree. There is completely no association between the phylogenetic tree of beta satDNA sequences and the phylogenetic tree of these species (Fig. 2), indicating that multiple horizontal gene transfers (HGT) have been taken place between species.

### Characterization of the diversity of beta satDNA sequences

To further compare the beta satellite sequences, a principal component analysis (PCA) was performed in order to identify putative groups without direct alignment. Visualization of sequences into the plane formed by the first two components of the PCA revealed five main groups, group ①-⑤, indicated by different colors in Fig. 3A. The GC contents of these five groups are 47.1%, 49.2%, 48.7%, 50.3% and 47.0%. The five groups were also marked out on the phylogenetic tree, showing that the groups can be largely distinguished from one another (Fig. 3B). We took out sequences from the five groups respectively, and performed alignments using Geneious 11, which showed that all the consensus sequences of the five groups have distinct characteristics at certain positions (Fig. 3C).

**Fig. 3.**
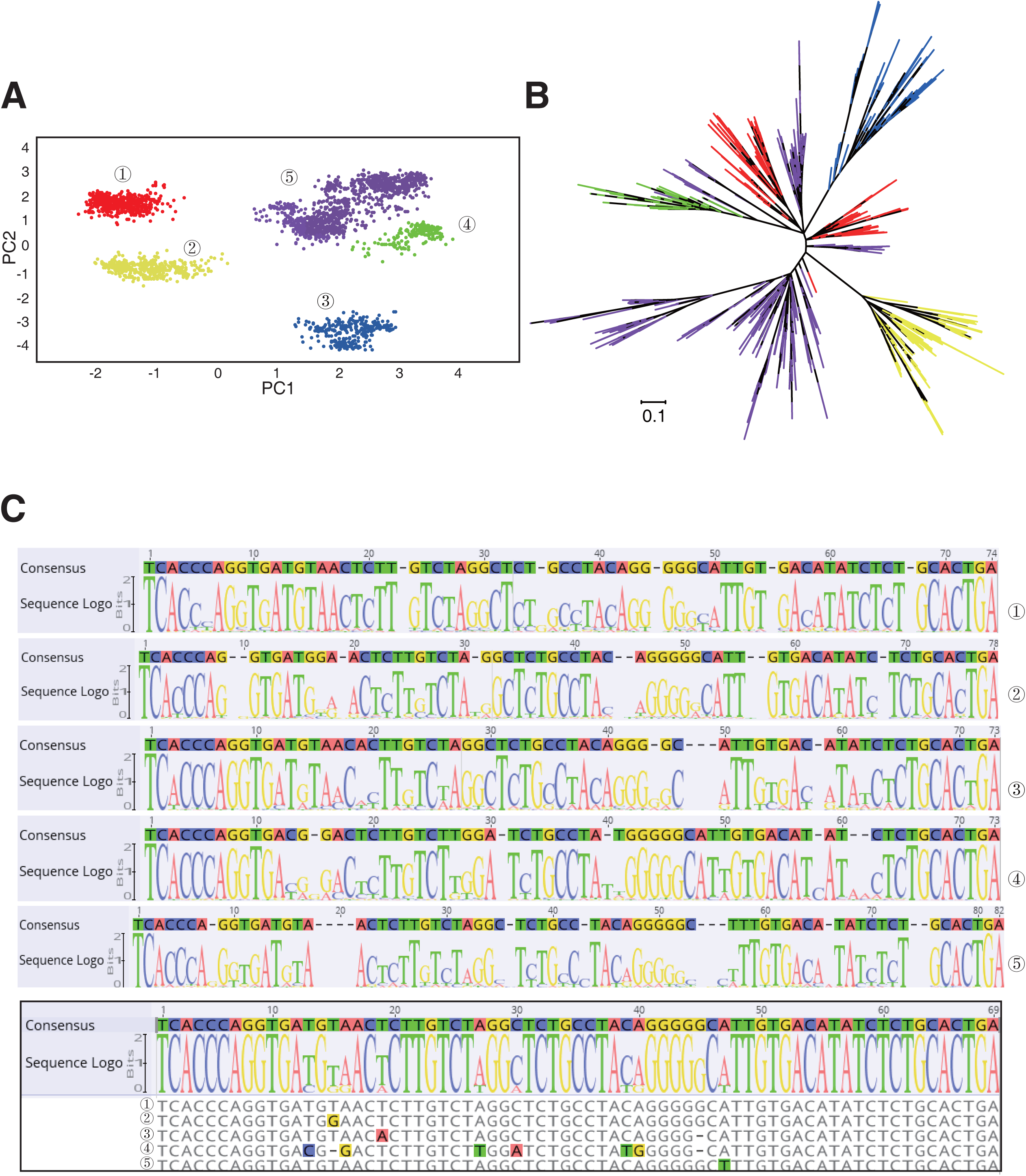
Characterization of beta satDNA diversity in 22 species. A, a PCA projection on principal components 1 and 2 for 1,274 sequences from 22 species. The sequences were divided into five groups (family ①-⑤), indicated by different colors. The PCA analysis was performed using Jalview. B, The distribution of different groups of beta satDNAs on phylogenetic tree. The different families were label by colors as described in A. C, The consensus sequences and sequence logos of the beta satDNA group ① - ⑤. The alignments were performed using Geneious 11.

## Discussion

The unequal exchange is a strong long-range ordering force which can keep tandem arrays homogeneous (Charlesworth et al., 1994). In this study, we found that beta satellite sequences have a high proportion of repeats within species (Table S1), indicating a certain degree of homogenization of these sequences. The homogenization process is much faster in the bisexual species, suggesting that meiotic recombination accelerates the homogenization process (Mantovani et al., 1997). Concerted evolution leads to higher homogeneity between the satDNA sequences of intraspecies than the sequences of interspecies (Dover, 1982; Kuhn et al., 2007). The higher-order alpha satDNAs are significantly more conservative within species than between primate species (Cacheux et al., 2016; Rudd et al., 2006). However, the beta satDNA sequences lack obvious conservative property within species (Fig. 2B), indicating the absence of complete concerted evolution, which might be related to a presumed reduction or suppression of meiotic recombination (Kuhn et al., 2007).

It is previously thought that beta satDNAs were originated in Catarrhini (apes and old world monkeys) and exploded in great apes (Cardone et al., 2004). However, deep sequencing allows us to detect low-abundance sequences that were hard to detect by traditional biochemical assays. We identified beta satDNA sequences in new world monkeys, prosimians, and even Dermoptera and Scandentia. Since recent studies placed Scandentia as a sister group of Glires (Meredith et al., 2011; Zhou et al., 2015), the origin of beta satDNAs could then be traced back to the common ancestor of Euarchontoglires. Of course, there is a concern that the copy numbers in new world monkeys, Prosimian, Dermoptera and Scandentia are low and even not found in the high-stringency dataset. However, a) the copy numbers in certain lesser apes and old world monkeys are also not high. It would not be too surprising that only a few copies of beta satDNAs were found in Prosimian and Dermoptera. b) All DNA samples have the risk of being contaminated by human DNA, whereas, beta satDNA sequences were not found in the genomic sequencing data of rodents or most other mammals. Therefore, we tend to believe that the existence of beta satDNAs in primitive primates is not an artifact. Taken together, the origin of beta satDNAs should be no less than the occurrence of the primate common ancestor (∼80 MYA), which is much earlier than people had thought.

Intriguingly, beta satDNAs were also found in some parasites of human and great apes. Both flatworms and nematodes are protostomes, while vertebrates (chordates) are deuterostomes. The separation of deuterostomes and protostomes was taken place more than 500 MYA, before the start of Cambrian. Therefore, the most reasonable explanation for this phenomenon is that these parasites acquired beta satDNAs from their primate hosts via HGT. Moreover, the observation that beta satDNAs don’t appear in other protostomes, such as insects, but are only found in the endoparasites of primates is another evidence for HGT.

The proportions of beta satDNAs in the genomes of *S. erinaceieuropaei* and *E. vermicularis* are even higher than those in orangutan genomes. And the patterns of phylogenetic tree of beta satellite sequences in these two parasites look more like the tree in human, which appears to have lots of recent branches, while the phylogenetic trees of beta satellite sequences in orangutans and old world monkeys look more primitive (Fig. 4). Thus, the current landscapes of beta satDNAs in parasites genomes might be the result of multiple HGT events between them and human or other members of Homininae, as well as the concerted evolution of beta satDNAs in the parasite genomes. The beta satDNA sequences within each species were also compared in pairs, and the identity distributions were shown in Fig. 5A. Similar distributions were found between beta satDNAs of *S. erinaceieuropaei* and human. Tandem repeats of beta satDNA units were identified from *S. erinaceieuropaei* genome contigs (Fig. 5B). The similar existences between satDNAs in the genomes of *S. erinaceieuropaei* and human suggest that beta satDNAs in the parasite genome may play similar role as they do in the hominid genomes.

**Fig. 4.**
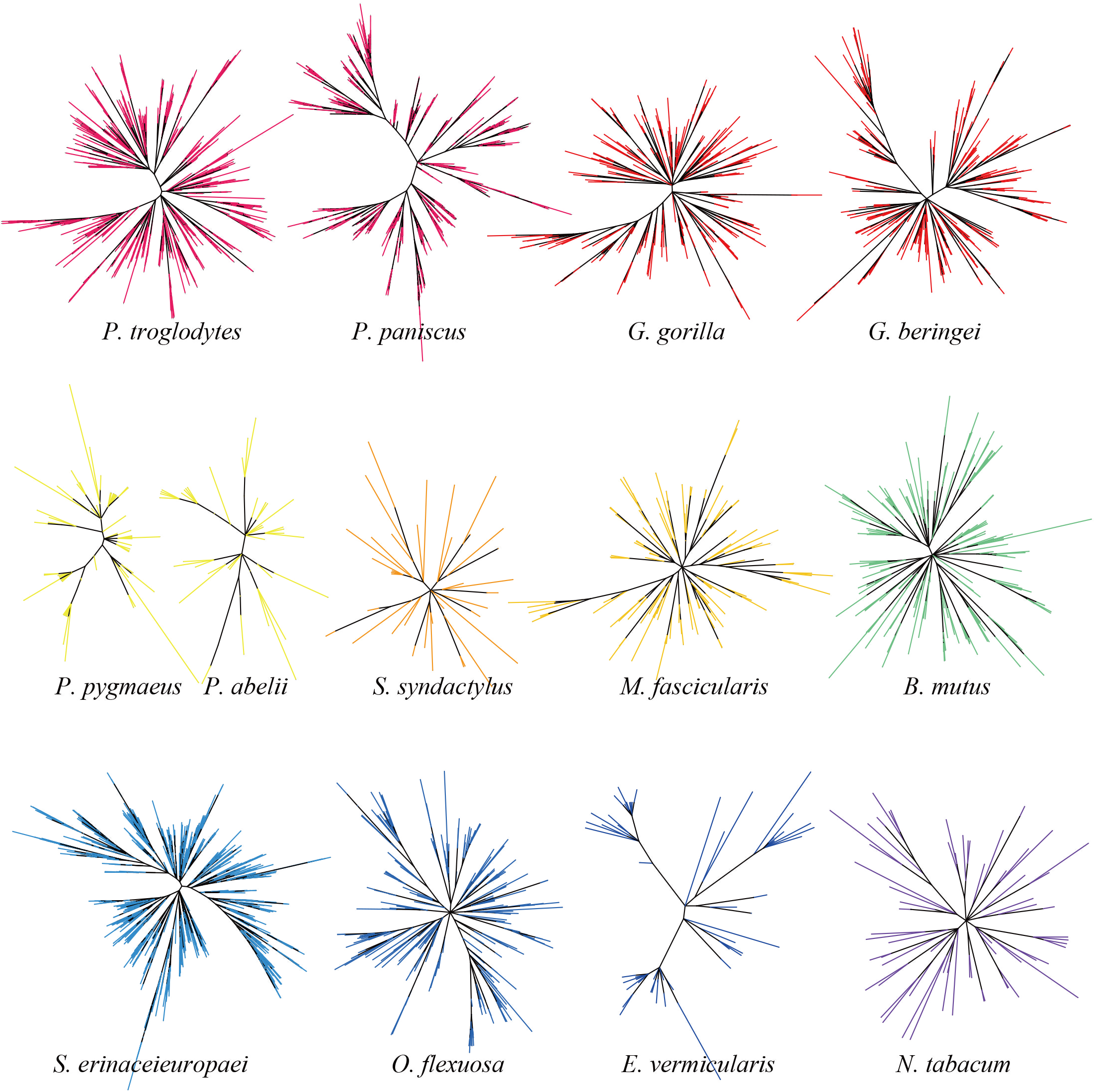
The phylogenetic trees of beta satDNA sequences from individual species. Trees were constructed using RAxML.

**Fig. 5.**
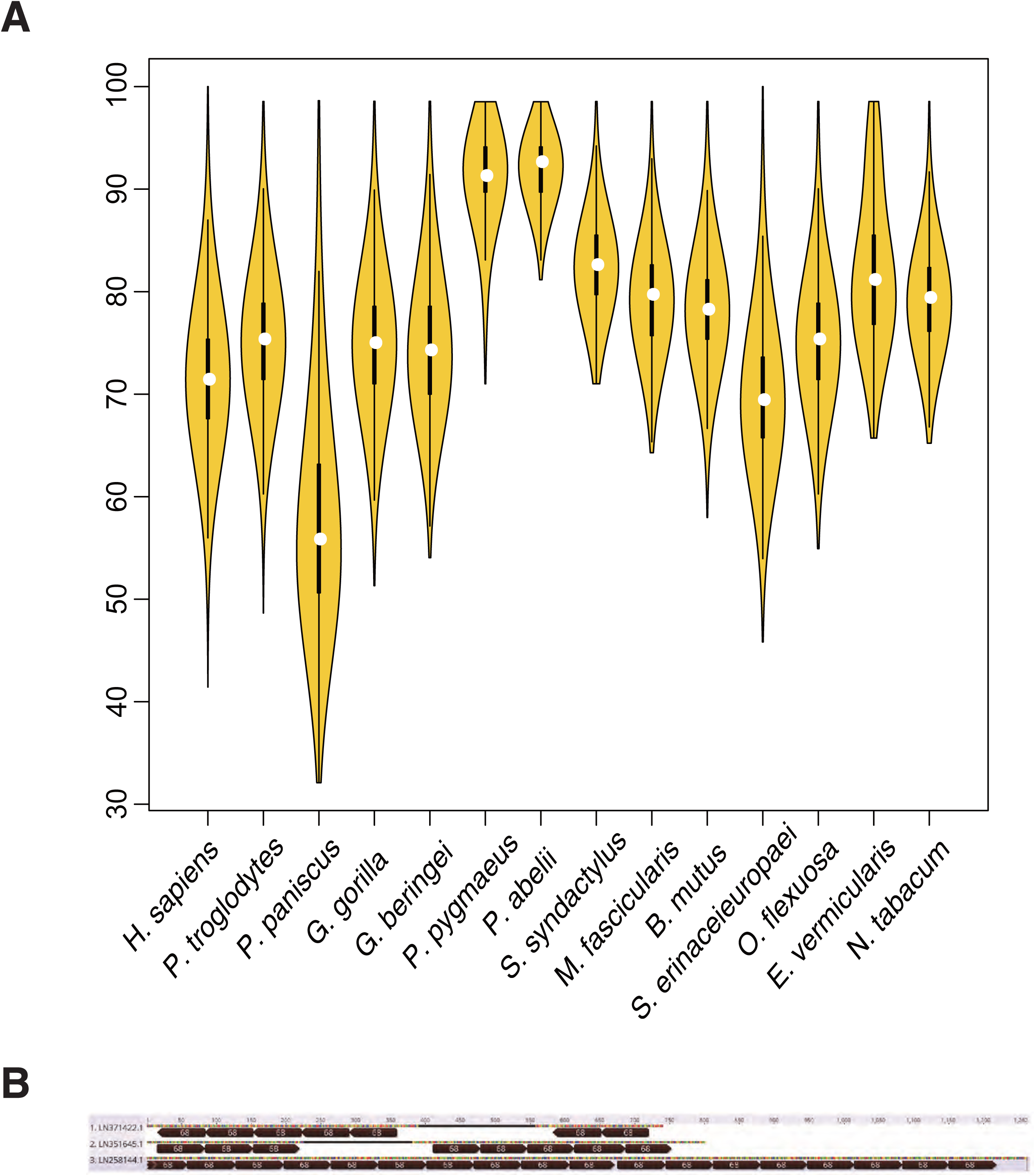
A, Tandem repeats of beta satDNA units on some contigs of the *Spirometra erinaceieuropaei* genome. B, The identity distribution of beta satDNA sequences within each of the species. Beta satDNA sequences within each of the species were compared in pairs and the distribution of the identities was shown by violin plot.

There are a number of copies of beta satellite sequences in the cattle genome. Since beta satDNA sequences were not found in other animals of Laurasiatheria (sheep, pig, dog or cat), the copies in cattle are more likely to be a result of HGT than inheritance from its mammal ancestor. Some parasites, such as *Onchocerca sp.* infect both human and cattle. They could be an intermediate for HGT from primates to cattle. We are so surprised that beta satDNAs were even found in a plant, *Nicotiana tabacum*. If the WGS data is reliable, the only explanation would be HGT. Blood-feeding insects and ticks are often chosen to account for HGTs between their animal hosts (Gilbert et al., 2010). Actually, some insects (e.g. certain species of Diptera) consume both animal blood and plant sap. They could have played the role of carrying DNA fragments from animals to plants.

Deep sequencing allows us to look into the highly repeated genomic sequences and find multiple HGTs of beta satDNAs. It was thought that HGT in eukaryotes are far less frequent than in prokaryotes. However, HGT in eukaryotes may be found far more common than had been thought as the sequencing technique becoming more and more powerful. Recent study showed that the transcripts of LINE-1 retrotransposon are essential for the mouse early embryo development (Jachowicz et al., 2017; Percharde et al., 2018). The reality is if you can’t get rid of certain exogenous sequences from your genome, you’d better collaborate with them, which is evolutionarily efficient and economical. Therefore, beta satDNAs in some non-primate genomes may also be functional in chromosome segregation and genome stability. It will not be surprising if beta satDNAs be identified from more non-primate genomes in the future.

## Materials and Methods

### Identification of beta satDNAs on human chromosome Y

The full sequence of the chromosome Y (NC_000024.10) was downloaded from National Center for Biotechnology Information (NCBI). The approximate locations of beta satDNAs on chrY were determined by making preliminary comments using Geneious 11 (Kearse et al., 2012), and the three regions containing beta satDNA copies, Ya, Yb and Yc were identified. Then the sequences of the three regions were mapped to a 68 bp beta satDNA reference sequence based on a 72 bp sliding window at 1 bp resolution (e.g. nt 1-72, nt 2-73…) using Geneious 11, and the dataset of the beta satDNA sequences on chrY was obtained. The scripts for extracting sequences were written in Python language.

### Identification of beta satDNAs in the genomes of different species

Using NCBI BLAST (nr/nt), beta satDNA sequences were found to distribute in at least 22 species. Since the satDNAs are highly repeated and difficult to be assembled into chromosome contigs, the next generation sequencing raw data were searched using BLASTn to collect as many beta satDNA sequences as possible. The raw sequencing data of the 22 species were downloaded from the NCBI SRA database (https://www.ncbi.nlm.nih.gov/sra) or BLASTed on line. Then the BLAST output files were filtered and the beta satDNA sequences were extracted using scripts written in Python. The percentages of beta satDNAs in each species were calculated using the following formula:

`number of hits / total number of reads in SRA databases * 100%`

### Phylogenetic analyses

Sequence alignment was performed using the CLUSTALW (Thompson et al., 1994), All phylogenetic analyses were conducted under maximum likelihood in RAxML (Stamatakis, 2014). The phylogenetic trees were colored using the online software ITOL(https://itol.embl.de/). Colored files were generated by a Python script.

### Principal component analysis (PCA)

Principal component analysis (PCA) for aligned DNA sequences were performed using Jalview (Waterhouse et al., 2009) to reduce the space complexity and enable data visualization on the first factorial plane. The input file was aligned using CLUSTALW.

### The identity distribution analysis

The data for the identity of sequences in pairs within each species were calculated by CLUSTALW and exported using Geneious 11. The violin plot was generated using an R package, vioplot.

## Supplementary materials

**Supplementary File 1** (see separated sequence file).

**Supplementary Table**

**Table S1.** Beta satDNA sequences in different species (*Same sequences that can be found in human sequencing data were excluded).

| Species | Total | Seqs with Ns<br>excluded | Same seqs<br>excluded | High-stringency* |
| --- | --- | --- | --- | --- |
| *Homo sapiens* | 201180 | 201134 | 22011 | 22011 |
| *Pan troglodytes* | 1036 | 1036 | 841 | 774 |
| *Pan paniscus* | 2018 | 2005 | 1211 | 1103 |
| *Gorilla gorilla gorilla* | 825 | 772 | 429 | 392 |
| *Gorilla beringei* | 3804 | 3802 | 437 | 398 |
| *Pongo pygmaeus* | 464 | 464 | 88 | 84 |
| *Pongo abelii* | 306 | 306 | 59 | 59 |
| *Symphalangus syndactylus* | 72 | 72 | 45 | 45 |
| *Hylobates moloch* | 17 | 17 | 9 | 8 |
| *Macaca fascicularis* | 203 | 203 | 144 | 31 |
| *Mandrillus leucophaeus* | 7 | 7 | 7 | 2 |
| *Saimiri boliviensis* | 4 | 4 | 4 | 0 |
| *Callithrix jacchus* | 1 | 1 | 1 | 0 |
| *Carlito syrichta* | 25 | 25 | 16 | 5 |
| *Prolemur simus* | 3 | 3 | 2 | 1 |
| *Galeopterus variegatus* | 12 | 12 | 4 | 0 |
| *Tupaia belangeri* | 1 | 1 | 1 | 0 |
| *Bos mutus* | 254 | 254 | 212 | 44 |
| *Spirometra erinaceieuropaei* | 4096 | 4091 | 2432 | 714 |
| *Onchocerca flexuosa* | 441 | 441 | 306 | 137 |
| *Enterobius vermicularis* | 369 | 369 | 65 | 23 |
| *Nicotiana tabacum* | 101 | 101 | 73 | 19 |

